# Estimating the Total Variance Explained by Whole-Brain Imaging for Zero-inflated Outcomes

**DOI:** 10.1101/2023.08.14.553270

**Authors:** Junting Ren, Robert Loughnan, Bohan Xu, Wesley K. Thompson, Chun Chieh Fan

## Abstract

Zero-inflated outcomes are very common in behavioral data, particularly for responses to psychological questionnaires. Modeling these challenging distributions is further exacerbated by the absence of established statistical models capable of characterizing total signals attributed to whole-brain imaging features, making the accurate assessment of brain-behavior relationships particularly formidable. Given this critical need, we have developed a novel variational Bayes algorithm that characterizes the total signal captured by whole-brain imaging features for zero-inflated outcomes . Our *zero-inflated variance* (ZIV) estimator robustly estimates the fraction of variance explained (FVE) and the proportion of non-null effects from large-scale imaging data. In simulations, ZIV outperformed other linear prediction algorithms. Applying ZIV to data from one of the largest neuroimaging studies, the Adolescent Brain Cognitive Development^SM^ (ABCD) Study, we found that whole-brain imaging features have a larger FVE for externalizing compared to internalizing behavior. We also demonstrate that the ZIV estimator, especially applied to focal sub-scales, can localize key neurocircuitry associated with human behavior.

## 1 Introduction

Non-invasively measuring the form and function of the human brain and mapping variation in these properties to cognitive and psychological outcomes has been fundamental for understanding the neural circuitry underpinning complex human behaviors.^1, 2, 3, 4, 5^ For example, magnetic resonance imaging (MRI) data are commonly used to identify brain features subserving specific neurocognitive functions^6, 7, 8^ and to build individual-level prediction models of specific behaviors.^9, 10^ Despite having been used in many published studies, two key problems in analyzing brain-behavior associations from MRI data have remained unsolved.

First, recent results of brain-behavior associations with large samples show that individual imaging features tend to have smaller effect sizes than previously expected, casting doubt on the validity and reliability of the “one-brain-feature-at-a-time” approach.^11, 12, 10, 13, 14^ In light of this, one approach is to estimate the theoretical upper bound of the variance explained given all imaging features from a given MRI modality.^15, 16^ However, the validity of these approaches for brain-behavior associations from MRI data remains to be examined. For example, it is unclear if the brain features for specific MRI modalities and behavioral outcomes are in fact not sparse.^16^ Moreover, the typical assumption of “ubiquity of effects” limits the ability of these variance components models to provide information about which features may be more or less important for driving brain-behavior associations.

Second, existing methodologies are built either for normally-distributed or binary outcomes.^16^ However, many behavioral outcomes are *semi-continuous*, i.e., characterized by a peak of values occurring at a minimum value along with typically right-skewed continuous values.^17, 18^ (Note, following the literature we refer to these distributions as being”zero inflated” even though the minimum may differ from zero.) Semi-continuous data arise frequently in applications, including medical costs,^19^ microbiome,^20^single cell gene expression^21, 22^ and psychological questionnaires.^23^

For example, the Child Behavior Checklist (CBCL), a widely used assessment of the mental state of children,^24, 25^ contains eight syndromal subscales and six DSM-oriented subscales, which yield right-skewed data with evident inflation at their minimum values, equal to fifty by scale construction^26, 3^ (Figure 1). Misspecifying semi-continuous outcomes as being normally-distributed, especially in highly zero-inflated and/or right-skewed data such as these, can cause severe, generally downward bias in estimates. Unfortunately, no analytical techniques so far are specifically developed to model the relationship between semi-continuous traits and brain imaging features.

**Fig. 1.**
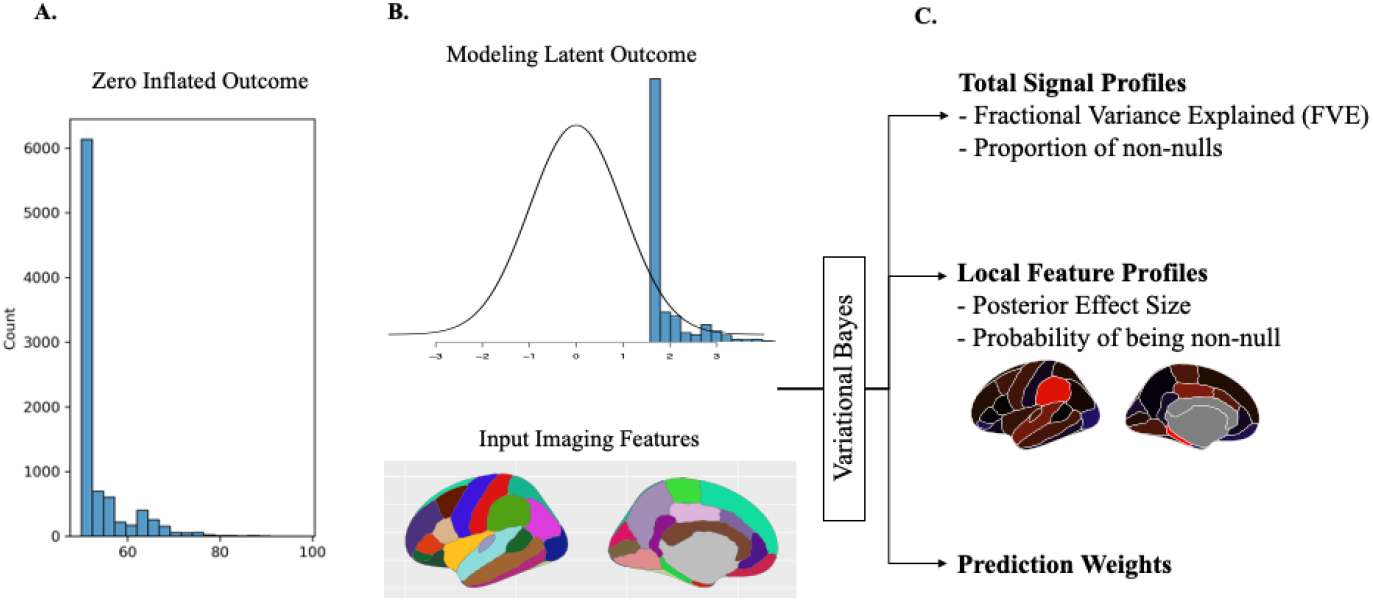
The schematic of the ZIV. A. The histogram of a zero-inflated outcome. B. ZIV assumes a latent outcome and the input imaging features have a linear relationship. C. Through variational Bayes algorithm, ZIV estimates both total signal profiles and the local feature characteristics simultaneously. The resulting posterior weights can be used for the prediction afterwards.

In light of the urgent need for methods that address these two issues, we present our newly developed Bayesian model, the zero-inflated variance estimator (ZIV). Figure 1 illustrates the design of ZIV. Given a set of whole brain imaging features, ZIV estimates the total variance explained by all features *en masse* (fraction of variance explained [FVE]; Figure 1 C, top section), proportion of non-null effects among all included features (Proportion of non-nulls), and identify which imaging features are most important for zero-inflated outcomes (Probability of being non-null; Figure 1 C, middle section). The posterior weights from the ZIV estimator can then can be used for prediction in an independent sample (Prediction Weights; Figure 1 C, bottom section). The practicality of the ZIV estimator is strengthened by our use of a variational inference algorithm, enabling inference with high-dimensional imaging data tractable. We demonstrate the validity and the utility of our method with comprehensive Monte Carlo simulations and empirical applications on a large-scale imaging cohort: the Adolescent Brain Cognitive Development^SM^ (ABCD) Study.

## 2 Method

### 2.1 Model Setup

The observed zero-inflated semi-continuous outcome *z*_*i*_ for subject *i* is modeled by positing a latent variable *y*_*i*_,

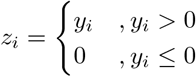

The latent variable and feature pairs 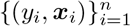 are assumed to follow a linear relationship:

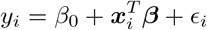

where the error terms *ϵ*_*i*_ are assumed to be independently and identically distributed as *N* (0, *σ*^2^), ***x***_*i*_ = (*x*_*i*1_, *x*_*i*2_, …, *x*_*ip*_) and ***β*** = (*β*_1_, *β*_2_, …, *β*_*p*_) are both *p*-dimensional column vectors.

We model the effects of the features as a mixture of priors from normally distributed non-nulls and point mass nulls (slab and spike prior):

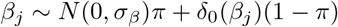

where *δ*_0_ is a point mass at 0. In addition, we assume the following prior for the global proportion of non-nulls, *π*, and the variance of the error terms, *σ*^2^, as:

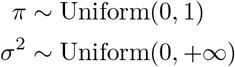

We denote by *θ* = (*σ*_*β*_, *β*_0_) the parameters optimized using gradient descent without a variational posterior. We reparameterize *β*_*j*_, *j* = 1, …, *p*, as

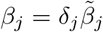

where

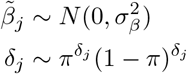

### 2.2 Variational Inference on the Posterior

We want to approximate the following true posterior

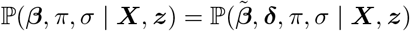

To do so we use the following variational distributions for approximation:

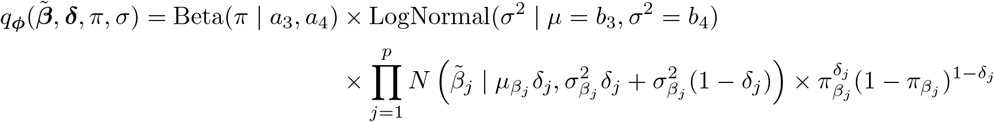

Notice that we have dependency between 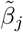 and *δ*_*j*_, where 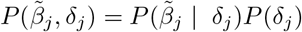.

We denote the variational parameters as ***ϕ*** = (*a*_3_, *a*_4_, *b*_3_, *b*_4_, ***μ***_*β*_, ***σ***_*β*_, ***π***_*β*_) for which we want to optimize over. To measure the distance between the true posterior and the approximate posterior, we use the Kullback-Leibler divergence (KL-divergence):

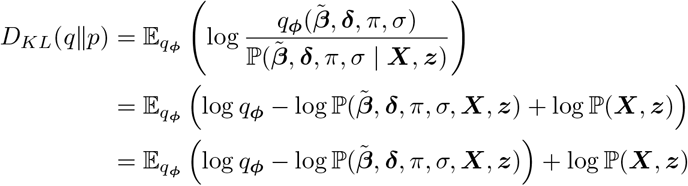

Therefore, we have

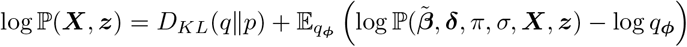

Since log ℙ (***X, z***) is a constant, in order to minimize *D*_*KL*_(*q*∥*p*), we maximize 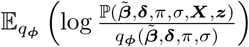 which is the Evidence Lower Bound (ELBO). For simplicity, we assume that the feature matrix ***X*** is fixed and always conditioned upon. The ELBO can be written in three separate parts:

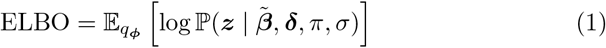

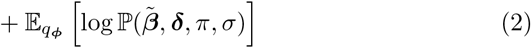

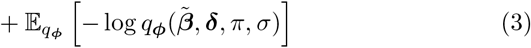

Equation 1 is the expectation of the data likelihood over the variational distributions, Equation 2 is the expectation of the prior likelihood and Equation 3 is the entropy of the variational approximation distribution. We use [27] for our stochastic gradient descend algorithm to minimize the negative of the ELBO. The details of the posterior inference, prediction and optimization methodologies are shown in the Supplemental Materials.

### 2.3 Model Inference

#### 2.3.1 Inferring Global Signal Architecture of Brain-Behavior Relationship

We used FVE and the proportion of non-nulls as the two key metrics to characterize the signal patterns of the brain-behavior associations as a whole. To estimate the FVE on the latent scale, we use the sampled values of latent linear effects in the posterior draw of the parameters. By definition, the variance captured by the latent linear outcome is

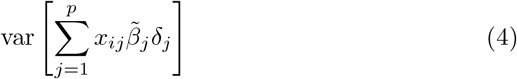

And we define the latent linear outcome value for each individual *i* at the current cycle is:

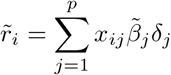

Then, the estimated variance in the current cycle is

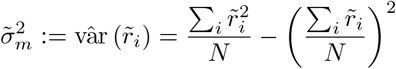

The estimated FVE on the latent scale is

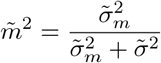

where 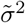 is the current estimate for the noise variance *σ*^2^. On the other hand, the inference for proportion of non-null effects can be done directly using the approximated posterior distribution Beta(*π* | *a*_3_, *a*_4_) for *π*.

#### 2.3.2 Localized Feature Inference and Selection

For individual feature selection, the model produces posterior distributions Bernoulli(*π*_*βj*_) for each feature *j*. Customized rules based on *π*_*βj*_ can be utilized to select out likely non-null features. In the simulation, we pick the top *k* highest *π*_*βj*_ as true associations, where *k* equal to the mean estimate of *π* multiply by the total number of features then divide by 2. There is a trade off between false discovery rate and sensitivity, so the rules should be created based on the actual needs of the researcher (Supplemental Materials). Given high degree of observed correlations between brain imaging features, ZIV provides an alternative but objective way to examine the feature importance.

#### 2.3.3 Prediction

The variational inference model approximates the posterior distributions of coefficient *j* using a mixture Guassian distribution 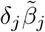 where 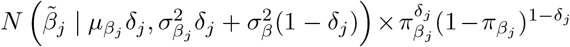. The variational parameters 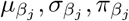 are optimized using gradient descent, and *δ*_*j*_ is sampled from the Bernoulli distribution with probability 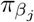. In order to obtain a point prediction, we directly take the mean of the coefficient posterior distributions and multiply them with the features as our predictions.

### 2.4 Monte Carlo Experiments

We performed Monte Carlo experiments to evaluate the performance of the ZIV model. In order to simulate synthetic data that closely resemble real data, we used actual task fMRI data for our imaging features while varying true model parameters to generate synthesized semi-continuous outcomes. The task fMRI data were sampled from the ABCD 4.0 Data Release, detailed in 2.5. We randomly assigned 0.5%, 1%, 5% or 10% of the features to have non-null effects. The non-null effects were generated from a normal distribution with mean 0 and standard deviation of 0.1. The latent outcome was set equal to the linear combination of the features multiplied by the corresponding coefficients adding normally-distributed noise. The observed outcome was truncated at 0 whenever the latent outcome was negative. The variance of the random Gaussian noise was determined by the empirical variance of the latent mean (linear combination of the features multiplied by the corresponding coefficients) and the pre-defined FVE parameter for simulation. We simulated data that have 0.05, 0.25, 0.5 or 0.8 fraction of variance explained. This resulted in a total of 16 different simulation scenarios (4 different percent non-null effects times 4 different pre-defined FVE’s). For each simulation setup, we generated 500 instances and aggregated over the estimates of the instances for final result.

First, we examined the performance of ZIV for inferring the global signal architectures, i.e. whether the FVE and the proportion of non-nulls covers the true values. For comparison, we also implemented a liability-based linear mixed effects model.^16, 28, 29^Estimation of mophemtricity from liability-based linear mixed effects model is one of the few existing high-dimensional imaging algorithms that focuses on characterizing the global signal.^16^ The liability based estimates were first proposed and implemented in the software, called GCTA,^28^ for human genetic studies and then applied to the neuroimaging data.^16, 29^ In contrast to ZIV, the GCTA model estimates the FVE, assuming the outcome is normally distributed and the signal architecture is ubiquitous across all input features.^28, 29^

Second, we investigated if ZIV improved the predictive performance by accounting for the zero-inflated distribution and non-null probabilities. We compared the predictive performance of ZIV with three other prediction models: ridge regression, LASSO, and the best linear unbiased predictor (BLUP) from the GCTA model. To evaluate the prediction performance of the models, we randomly split the data into 80% training and 20% testing sets for each outcome-modality pair. The performance metric we used was the mean absolute error (MAE). Compared to mean-squared error (MSE), MAE is less sensitive to outliers that are abundant in non-Gaussian data^30^ and hence more appropriate for semi-continuous, highly skewed data. In addition, MAE is more interpretable than MSE because it is assessed in the same units as the data, while the MSE is in squared units.^31^ Finally, we compared the computational speed and accuracy between variational inference and the traditional Markov chain Monte Carlo (MCMC) algorithm using a similar simulation setup.

### 2.5 Empirical Application

We investigated the relationship between the psychopathology and Region of Interest (ROI) brain measurements, one imaging modality at a time, using data from the Adolescent Brain Cognitive Development (ABCD) Study. ABCD is the largest study of neurodevelopment in the United States. N=11,880 youth aged 9-11 at baseline were recrutied from 21 different sites around the country; they are currently undergoing annual in-person evaluations for over a decade by the end of the study.^32, 33^ Data are released publicly on an annual basis via the National Institute of Mental Health Data Archive (NDA, https://data-archive.nimh.nih.gov/abcd). More details about the study can be found at https://abcdstudy.org/. In this application, we used data from the ABCD 4.0 National Data Archive release (NDAR DOI:10.15154/1523041).

#### 2.5.1 Child Behavior Checklist scores

The Child Behavior Checklist (CBCL) is a widely-used tool utilized for evaluating an extensive range of emotional and behavioral problems in children.^34, 35, 36^ It uses a scoring system where responses are labeled as 0 (not applicable), 1 (partially or occasionally applicable), or 2 (completely or frequently applicable). The CBCL is comprised of 113 items that measure aspects of the child’s behavior across the past six months. The CBCL provides a total score, along with scores on eight syndrome subscales and six subscales oriented around the Diagnostic and Statistical Manual of Mental Disorders (DSM). The eight syndrome subscales include: (1) anxiety/depression; (2) social withdrawal/depression; (3) somatic complaints; (4) issues with social interaction; (5) thought disturbances; (6) attention issues; (7) rule breaking; and (8) aggressive behavior. Subscales from these eight syndromes can be further summarized into three problem scales: internalizing problems (comprising anxiety/depression, social withdrawal/depression, physical complaints), externalizing problems (comprising violations of rules, hostile behavior), and total problems. The six DSM-oriented scales encompass: (1) depressive disorders; (2) anxiety disorders; (3) somatic disorders; (4) attention-deficit/hyperactivity disorders; (5) oppositional defiant disorders; and (6) conduct disorders. Scores obtained from the CBCL are usually highly right skewed.^37^ In particular, the t-standardized score, a preferred scoring that aims to reduce over-interpretation, exacerbates the violation of normal assumption by left truncation of the raw score at 50, leading to inflation at this minimum value.^38^ Here, we focus on the t-standardized scores of all eight syndrome scales, six DSM-oriented scales, and three summary problem scales, investigating how brain features associate with these semi-continuous scores.

#### 2.5.2 Multimodal Imaging Measures

Neuroimaging data were consolidated across all 21 data collection sites and processed by the ABCD Data Analysis Informatics and Resource Center and the ABCD Image Acquisition Workgroup.^39^ Data were then obtained at the region of interest (ROI) level for the five MRI modalities available in the ABCD data release: 1) structural T1 MRI (sMRI), which measures cortical and subcortical morphometry; 2) diffusion tensor images (DTI), which are sensitive to the fiber structures of human brain; 3) restricted spectrum images (RSI), which summarize the properties of tissue compartments; 4) task functional MRI (task fMRI), consisting of event-related contrasts capturing change in the fMRI signal in response to stimuli;^40^ and 5) resting state functional MRI (rsMRI), consisting of connectivity measures across Gordon parcellations and subcortical regions, partitioned into modular networks.^41^

The number of features per modality were as follows: 1) *p* = 1,186 measures from sMRI; 2) *p* = 2,376 measures from DTI; 3) *p* = 1,140 from RSI; 4) *p* = 885 from the three fMRI tasks; and 5) *p* = 416 from rsMRI. For all imaging modalities except rsMRI, ROIs were restricted to the Desikan cortical atlas.^42^ As described above, rsMRI was based on a modular network partition. Both DTI and RSI features included metrics from segmented major fiber bundles, in addition to the cortical and subcortical ROIs. Casey et al. (2018)^40^ offers in-depth information about the imaging acquisition, processing procedures, and quality control metrics in the the ABCD imaging data. All features were standardized to have zero mean and unit standard deviation before entering them into the ZIV models.

#### 2.5.3 Participant Inclusion

For each outcome (CBCL scores) and image modality pair, we excluded participants who were missing outcome observations or lacked the corresponding modality ROI measurements. We randomly sampled one member from each family if there were multiple siblings within a family. Within each participant, a single observation was randomly sampled if that participant had multiple MRI assessments across visits. The number of observations included for each image modality for all outcomes is shown in Supplementary Materials Table 7.3. In all ZIV models we adjusted for race, age, MRI scanner serial info and software versions, and sex at birth as potential confounders. Variance due to confounders is thus not included in the calculation of FVE for brain imaging features.

## 3 Results

### 3.1 Monte Carlo Results

Simulation results are shown in Table 1. The ZIV model provides unbiased estimates of the FVE’s under all scenarios (Table 1); in contrast, GCTA severely underestimates them in all scenarios. Furthermore, unlike GCTA, our ZIV model provides estimates for the proportion of non-nulls. The Credible Interval (CI) for FVE and proportion of non-nulls provide good coverage of the true values, with less than 5% error rates (Table 1). In the context of prediction performance, the ZIV model out-performed predictions made by GCTA, LASSO and Ridge (Table 1). On average across all the scenarios, ZIV has a 49.7%, 19.8%, and 21.2% improvement over GCTA, LASSO, and Ridge in terms of MAE, respectively. Further detail of the simulations regarding inference for localized non-null features and the comparison between variational inference and MCMC can be found in the Supplementary Materials.

**Table 1.** Simulations with various brain-behavior signal architectures.

| True FVE<br>True Non-null Proportion | 0.25 |  |  |  | 0.50 |  |  |  | 0.80 |  |  |  |
| --- | --- | --- | --- | --- | --- | --- | --- | --- | --- | --- | --- | --- |
|  | 0.005 | 0.010 | 0.050 | 0.100 | 0.005 | 0.010 | 0.050 | 0.100 | 0.005 | 0.010 | 0.050 | 0.100 |
| **Point Estimates** |  |  |  |  |  |  |  |  |  |  |  |  |
| FVE by GCTA | 0.124 | 0.127 | 0.129 | 0.131 | 0.260 | 0.263 | 0.265 | 0.269 | 0.438 | 0.447 | 0.449 | 0.450 |
| FVE by ZIV | 0.241 | 0.241 | 0.236 | 0.230 | 0.487 | 0.487 | 0.479 | 0.472 | 0.792 | 0.791 | 0.786 | 0.780 |
| Inferred proportion of non-nulls by ZIV | 0.011 | 0.016 | 0.040 | 0.053 | 0.010 | 0.015 | 0.044 | 0.065 | 0.008 | 0.014 | 0.049 | 0.078 |
| **95% CI Coverage Error Rate** |  |  |  |  |  |  |  |  |  |  |  |  |
| FVE ZIV Error Rate | 0.010 | 0.002 | 0.000 | 0.008 | 0.010 | 0.002 | 0.004 | 0.000 | 0.076 | 0.028 | 0.004 | 0.002 |
| Error Rate of proportion of non-nulls | 0.000 | 0.000 | 0.000 | 0.010 | 0.000 | 0.000 | 0.000 | 0.000 | 0.000 | 0.000 | 0.000 | 0.000 |
| **Prediction - Mean Absolute Errors (MAE)** |  |  |  |  |  |  |  |  |  |  |  |  |
| ZIV | 0.397 | 0.593 | 1.468 | 2.067 | 0.231 | 0.342 | 0.850 | 1.213 | 0.115 | 0.169 | 0.423 | 0.615 |
| GCTA | 0.507 | 0.759 | 1.855 | 2.585 | 0.448 | 0.663 | 1.599 | 2.265 | 0.620 | 0.888 | 2.197 | 2.930 |
| LASSO | 0.464 | 0.693 | 1.708 | 2.402 | 0.280 | 0.413 | 1.016 | 1.435 | 0.163 | 0.238 | 0.591 | 0.842 |
| Ridge | 0.476 | 0.710 | 1.729 | 2.411 | 0.287 | 0.423 | 1.034 | 1.452 | 0.167 | 0.244 | 0.599 | 0.849 |

**Table 2.** Number of observations and features for data application.

| modality | N | p |
| --- | --- | --- |
| Resting state MRI | 9501 | 416 |
| Task functional MRI | 8893 | 885 |
| Structural MRI | 9426 | 1186 |
| Diffusion tensor images | 9454 | 2367 |
| Restricted spectrum images | 9455 | 1140 |

### 3.2 ABCD Study Results

We used ZIV to infer the signal architecture of the CBCL subscales, one imaging modality at a time. Results are summarized in Figure 2A. Across CBCL scales, the estimated FVE ranges from 0.6% to 4.9%. Imaging modalities have a similar range of FVE’s given the same CBCL subscale, with the highest magnitude of FVE in the DSM-oriented Conduct subscale and the Rule Break syndrome subscale. However, the proportion of non-nulls differs across imaging features. Across CBCL subscales, rsfMRI has the highest proportion of non-null compared to other imaging modalities, despite having a similar range of FVE’s.

**Fig. 2.**
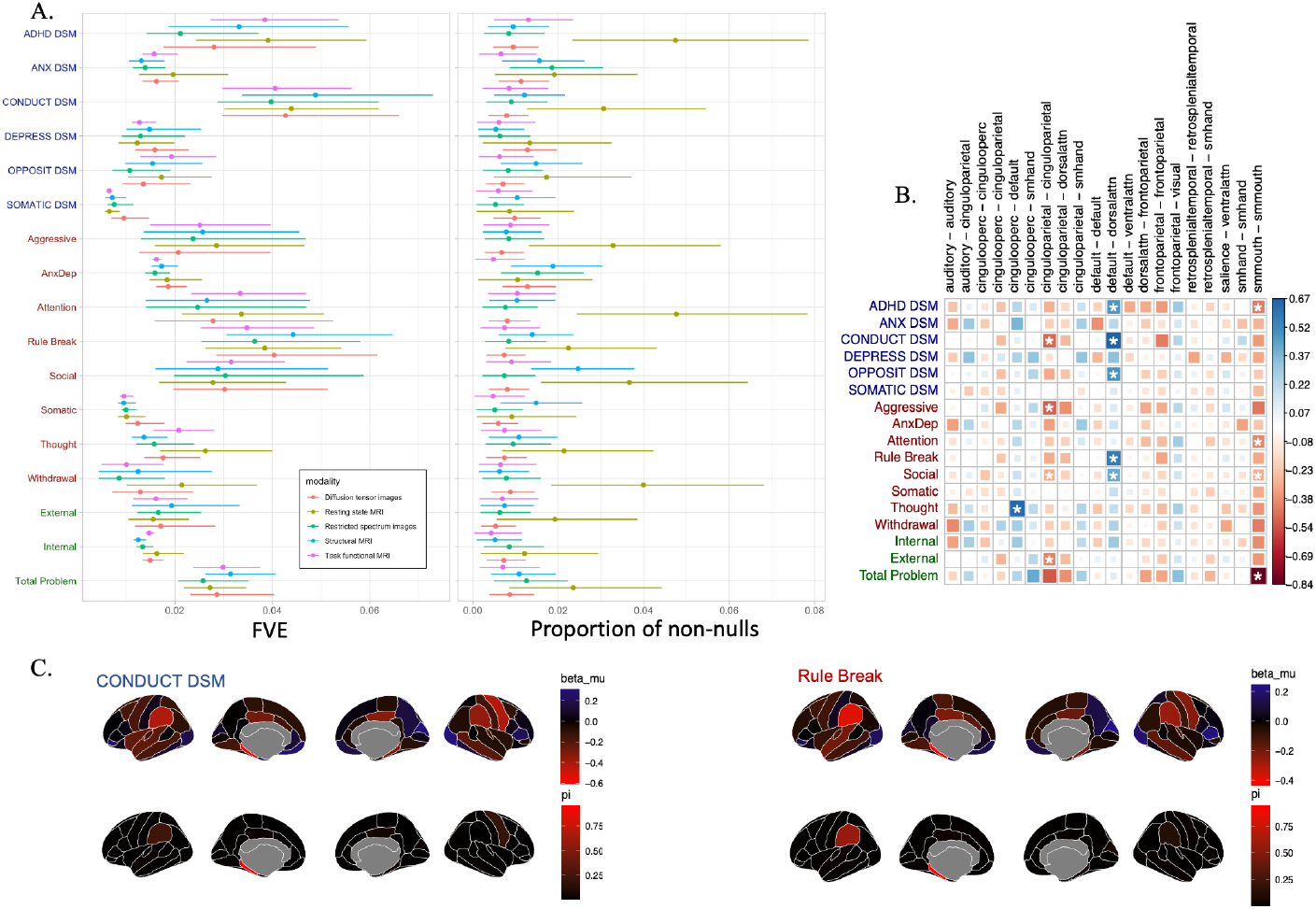
Brain and CBCL behavior associations in ABCD. A. The global signal architectures of CBCL across imaging modalities. B. Posterior estimates of the local features of rsfMRI. The coloring and size of the square in a cell represent the posterior effect size of each measure pair while the asterisk marks indicate the probability of being non-null exceeding 50%. C. Posterior estimates of the local features of sMRI for two CBCL scales that have highest FVE. The posterior effect sizes are illustrated on the upper row and the posterior probabilities are illustrated on the lower row.

The estimated proportion of non-nulls are all less than 5% of included features. Posterior estimates thus indicate brain-behavior signals are concentrated on smaller subsets of ROIs with an overall background of weaker effects (Figure 2 B and C). Between-network connectivity between the default mode and dorsal lateral attention networks is associated with CBCL scales in three DSM-oriented scales (ADHD, Conduct, and Opposition) and two syndrome scales (Rule break and Social). Within cingulo-parietal network connectivity, on the other hand, is found to be most closely related to externalizing behaviors. The somatosensory network has strong associations with all CBCL subscales, although most prominently in the ADHD, Attention, Social, and Total Problem subscales (Figure 2 B).

Mirroring results from rsfMRI, local signals from the associations with sMRI measures were more evident in the cingulate, pariental, and somatosensory regions, as shown in the Figure 2 C. The volume of the parahippocampus was inferred to have the highest probability of being non-null compared to all other regions.

## 4 Discussion

We demonstrated that the ZIV model can efficiently infer the total global signal while simultaneously localizing the driving features, given zero-inflated outcomes and high-dimensional imaging predictive features. In simulations using realistic high-dimensional features from task fMRI, ZIV out-performed other tools in estimating the true values of FVE and proportion of non-null signal, as well as in predictive accuracy on independent testing sets. By adopting variational inference, ZIV provides posterior estimates within minutes using a modern laptop computer. ZIV is hence useful and practical for application to large-scale brain-behavior analyses of many existing datasets.

Analyzing zero-inflated outcomes without considering the violation of normality assumption can lead to serious bias. As showcased in our simulations, FVE estimated by methods like GCTA are severely downward biased. ZIV formally models the zero-inflated outcome as a result of truncation of a partially observed latent variable, hence more accurately capturing the global signal architecture. With correct model specification, ZIV also benefits during prediction, performing better than GCTA, LASSO, and Ridge regression in all settings.

Our empirical findings of applying ZIV to the ABCD cohort indicate that, regardless of imaging modality, relevant brain features are most consistently detected for externalizing symptoms. Conduct problems, rule breaking behaviors, and total problem scales exhibited the highest FVE across measures. It is perhaps unsurprizing that externalizing symptoms were most strongly linked with brain morphology and function, with ADHD and conduct problems forming some of the most prevalent mental disorders in early adolescence.^43^ Indeed, recent analysis within the ABCD sample found these behaviors to be most strongly predicted by genetics.^44^ Taken together these results indicate that externalizing symptom assessments may exhibit more variability in this young adolescent sample leading to a greater ability to detect associations. A recent high impact paper used ABCD data to argue that the strength of brain-behavior associations were much smaller than previously thought.^45^ In this work researchers presented cross-validation predictions claiming that rsfMRI features explain around 1 percent of variance of the CBCL Total Problem while much less for the CBCL internalizing and externalizing measures. We find this previous work likely underestimated proportion of variance explained, in part due to a mispecified model assuming normality of variables. the current work tackles this problem directly and in so doing provides a more comprehensive picture of associations between brain and adolescent mental health.

In particular, our estimates of the proportion of non-null effects indicate that, for the item-level behavioral measures among youth, brain-behavior signals are not ubiquitous across brain regions. This is in contrast with reports that focus on more complex and normally distributed outcomes, such as intelligence scores.^46^ This sparseness also violates assumptions used in methods such as GCTA,^28, 29, 16^ rendering them inappropriate to model such effects. The sparseness of the non-null effects across brain regions enables analyses to partition out neurocircuitry related to components of behavior-such as those captured by item-level measures. Here we have showcased that ZIV is well suited for this purpose since many of the item-level behavioral measures are zero-inflated and heavily skewed.

Our analyses domonstrated that default mode network connectivity with the dorsal lateral attention network has consistent associations with ADHD, Conduct, Opposition, Rule breaking, and Social scales. The mis-engagement of the default mode network with the attention network has been posited as a key driver for attention issues among youth.^47^ Concordant with the rsfMRI results, our analysis on the volumetric measures from sMRI show that the reduced volumes in parietal, cingulate, and parahippocampal regions are consistently associated with Conduct and Rule breaking scales. These results suggest that circuitry linking these regions is more salient to externally-manifested behavior, rather than attention alone.

In sum, our development and efficient implementation of the ZIV mddel provides a necessary tool to investigate brain-behavioral relationships zero-inflated, highly-non-normal data. In these cases, ZIV produces unbiased estimates of the global signal from high-dimensional imaging data while providing detail on signal localization. Because of the high prevalence of semi-continuous data in many fields, ZIV could be applied to analyses beyond brain-behavior research.

## 5 Code Availability

The code associated with this research is available on GitHub at https://github.com/junting-ren/ZIV. This repository contains all necessary code and instructions to replicate the analyses described in this paper.

## 6 Acknowledgement

This work was supported by grant R01MH122688, RF1MH120025, and R01MH128959 funded by the National Institute for Mental Health (NIMH). The ABCD Study is supported by the National Institutes of Health and additional federal partners under award numbers: U01DA041022, U01DA041028, U01DA041048, U01DA041089, U01DA041106, U01DA041117, U01DA041120, U01DA041134, U01DA041148, U01DA041156, U01DA041174, U24DA041123, and U24DA041147. A full list of supporters is available at https://abcdstudy.org/federal-partners.html. A listing of participating sites and a complete listing of the study investigators can be found at https://abcdstudy.org/consortiummembers/. ABCD consortium investigators designed and implemented the study and/or provided data but did not necessarily participate in the analysis or writing of this report. This manuscript reflects the views of the authors and may not reflect the opinions or views of the NIH or ABCD consortium investigators.

## 7 Supplement

### 7.0.1 Expectation of the data likelihood

Due to the truncation of the observed phenotype *z*, the log data likelihood for *z* = 0 is

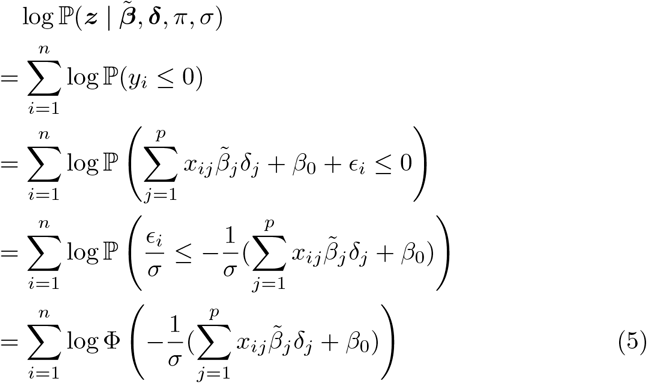

where Φ is the cumulative distribution function for standard normal distribution. The log data likelihood when *z >* 0 is given by

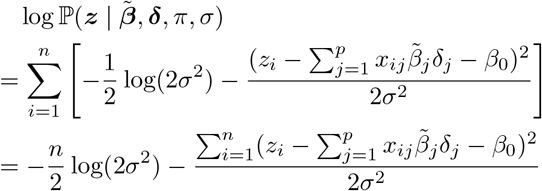

The variational parameters that need to be taken expectation of are 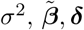. When *z* = 0, we directly plug in the current estimate 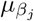 in place of 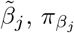 in place of *δ*_*j*_, and 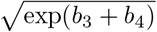 (mode of the standard deviation variational distribution) in place of *σ* into Equation 5 as approximation to the expectation of Equation 5.

For the expectation of the log data likelihood when *z >* 0, it is possible to calculate the exact value. Due to the factorization of the approximation distribution, we can first take the expectation with respect to *σ*,

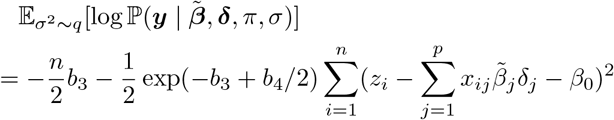

As for 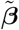, ***δ***, we need to expand 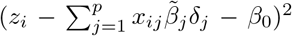 before taking the expectation. We use the Gumbel-softmax [48] to sample *δ*_*j*_ and reparametrization trick [49] to sample 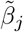 so that there will be gradients for the respective parameters using Monte Carlo integration. The best temperature for Gumbel-softmax is 1. During simulation experimentation for the Monte Carlo integration model, it was observe that even by taking more samples for the integration, the performance did not improve. From further investigation, this is because simply by plugging in 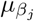 and 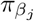 in place of *β*_*j*_ and *δ*_*j*_, the only difference between the true expectation and the plug-in approximation is on the quadratic term, where the true expectation contains 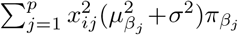 comparing the approximation 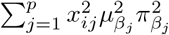 for each *i*. By taking the difference, we can obtain the true expectation. The true expectation of the term 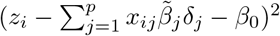 is:

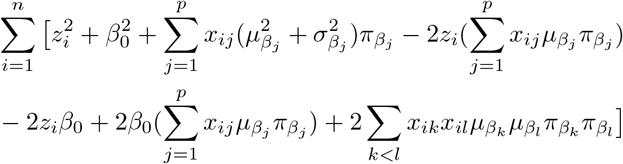

Note that this is exactly the same as the variational inference posterior if *β*_*j*_ and *δ*_*j*_ are independent since the conditional dependency is not introduced in the data likelihood. Therefore, in the final algorithm, we used the true expectation of the data likelihood to optimize over instead of the Gumbel-softmax approximation version.

### 7.0.2 Expectation of the prior likelihood

Expanding the inside of equation 2,

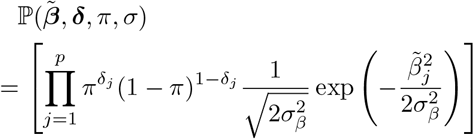

Therefore, taking the log, we have

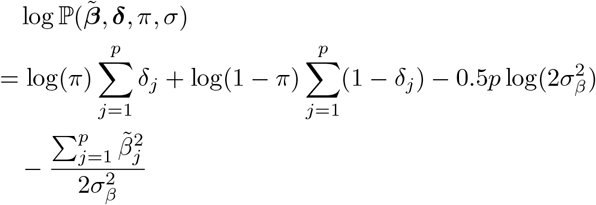

Now, we know

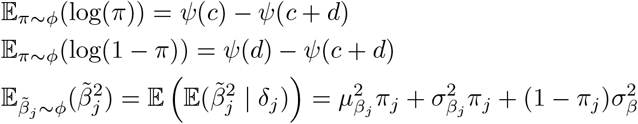

where *ψ* is the digamma function. From linearity of the expectation, we can obtain

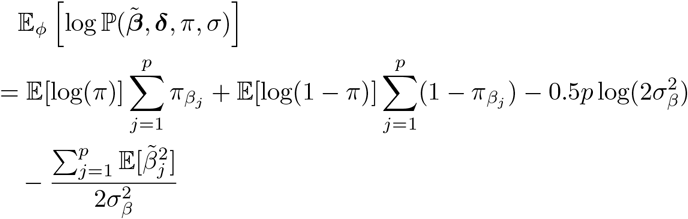

### 7.0.3 Entropy

Since the variational approximation distributions can be factorized, we con-sider the entropy for 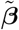, ***δ*** first:

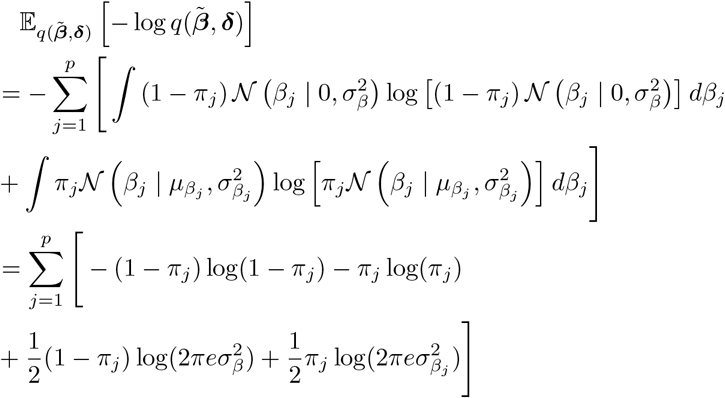

The entropy for *π* Is

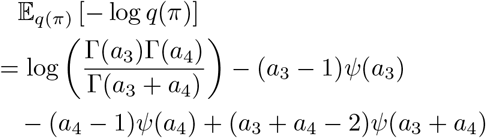

where Γ is the gamma function. The entropy for *σ* Is

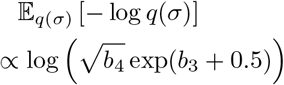

#### 7.1 Model Optimization

The prior hyper-parameters for global proportion of non-nulls *π* and error variance *σ*^2^ are pre-defined as shown in Section 2. The prior variance of coefficients *σ*_*β*_ and the intercept *β*_0_ are optimized using gradient descent similar to the variational parameters.

Majority of the parameters that needs to be optimized are initialized randomly using either standard Gaussian distribution or Uniform distribution.^50^ For the posterior approximating Beta distribution for *π*, the parameters *a*_3_ and *a*_4_ are initialized with 1.1. For the posterior approximating LogNormal distribution for *σ*^2^, the parameters *b*_3_ and *b*_4_ are initialized at 10.0 and 0.1, respectively.

We used the Adam optimizer^27^ with learning rate of 0.5, with betas equal to (0.9, 0.999). An exponential learning rate scheduler is used with decay multiplicative factor 0.8 for every 1000 epochs. Early stop is implemented whenever the difference between the current loss and minimum loss is less than 1% for 200 epochs, with a max of total training epochs of 20000. Bayesian variational inference inherently guards against overfitting, so there is no need for a validation set to be used for early stopping. In this context, early stopping has been implemented specifically to expedite the training process.

#### 7.2 Supplemental Experimentation Result

For the main simulation, we pick the top *k* highest 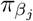 as causal, where *k* equal to the mean estimate of *π* multiply by the total number of features then divide by 2. The sensitivity and False-discover rate (FDR) for different proportion of non-null features are shown in Figure 3. We can see that using this feature selection approach that focus more on controlling false discoveries, we achieve a low FDR of at most 0.25. As the FVE increases, the FDR decreases and can be as low as 0. The sensitivity increases with FVE and lowers with the true proportion of non-null features. This is because as the percentage of causal features increases, features with high correlations with casual features may be erroneously selected.

**Fig. 3.**
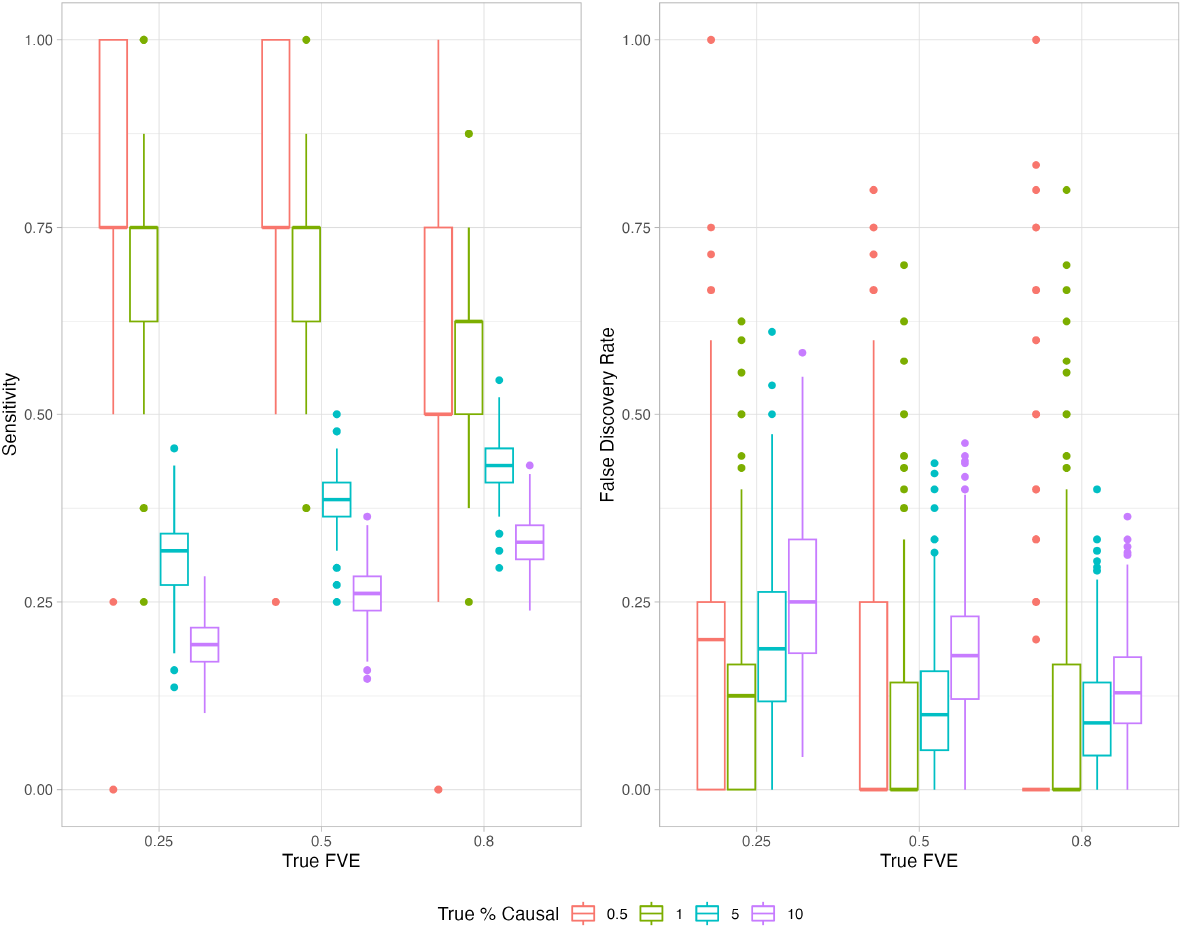
Sensitivity and FDR for causal feature selection

In addition, we compared the model inference speed and estimation accuracy between VI and Markov chain Monte Carlo (MCMC). Due to the high computation requirement of MCMC, we used the same simulation setup but limiting the number of observations to 5000, number of features varying from 100 to 500, FVE equals to 0.5 and percentage of causal equal to 10%. For each comparison setup, a total of 100 instances are generated. The result is shown in Figure 4. As the number of features increases, the fitting time for MCMC increases exponentially, whereas the time increases linearly for VI. The FVE for both approaches are estimated with high accuracy achieving unbiased estimates at 0.50. The VI inference approach achieves smaller variance comparing to MCMC under this simulation setup.

**Fig. 4.**
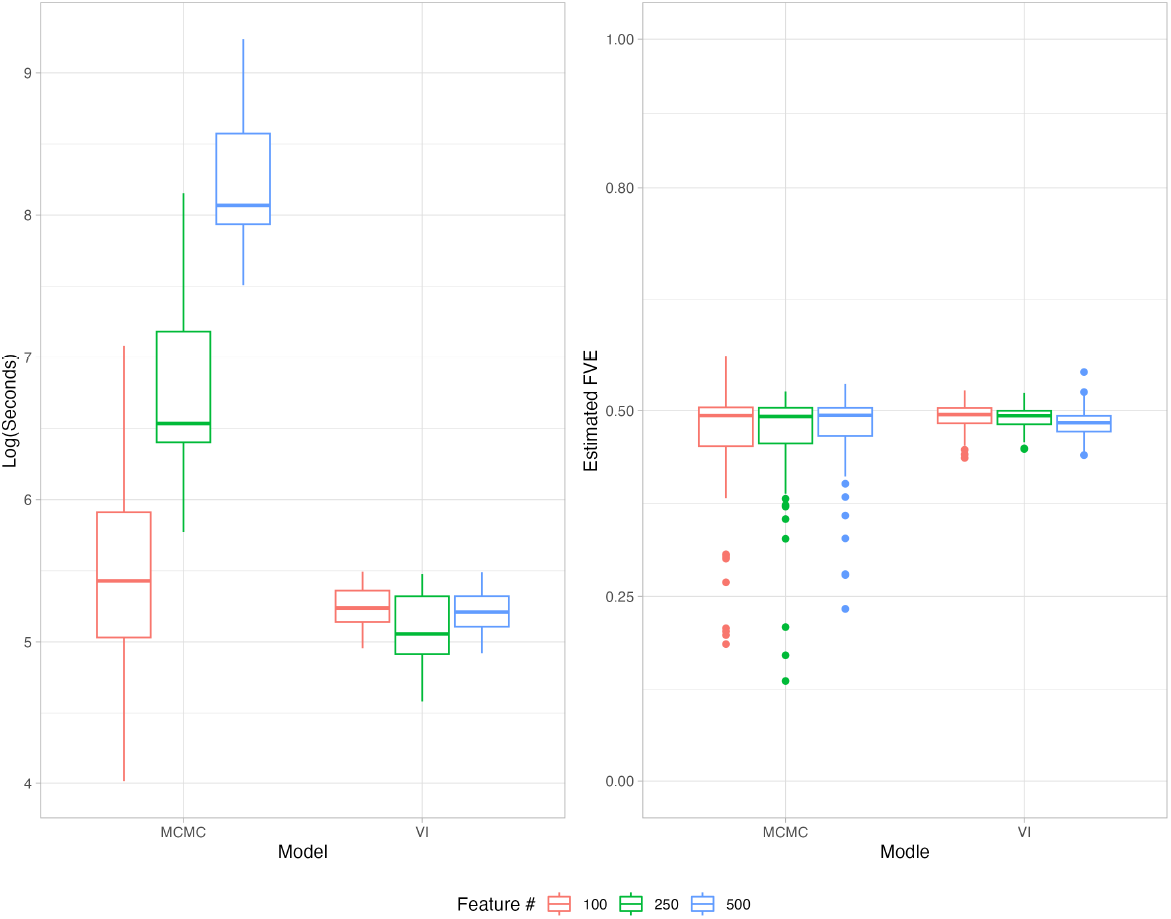
Comparing VI to MCMC.

#### 7.3 Data Application Sample Size and Feature Number

